# Tis But a Scratch! Negligible fitness costs of AalDV2 infection in *Aedes albopictus* under fluctuating temperatures and implications for viral biocontrol

**DOI:** 10.64898/2026.07.06.736729

**Authors:** N Sacco, M Perriat-Sanguinet, P Makoundou, A M’Sakni, M van Munster, C Boëte

## Abstract

*Aedes albopictus* is a major arboviral vector whose global expansion, driven by human activities and climate change, poses a growing public health concern for a number of neglected tropical diseases in both tropical and temperate regions. As a poikilotherm, its biology and population dynamics are strongly influenced by temperature, thereby shaping disease transmission. To thwart and control its geographic expansion, effective vector control strategies are increasingly critical.

Densoviruses (DVs), such as AalDV2, are being explored as mosquito viral biocontrol agents due to their restricted host range and ability to disseminate through oviposition sites. However, the influence of environmental parameters on the interactions between *Ae. albopictus* and AalDV2 remains poorly understood. This makes their performance under realistic, fluctuating thermal regimes difficult to estimate.

In this study, we investigated the combined effects of temperature and AalDV2 exposure on *Ae. albopictus* survival and development across its full life cycle. Mosquitoes were reared under fluctuating temperature regimes (26-28°C and 32-34°C, 12:12 day–night cycles) and exposed to AalDV2 or a control treatment.

Chronic exposure to 32–34°C significantly reduced overall survival, decreasing median lifespan by approximately 10 days (HR=2.21, p=0.0018), with a deleterious effect increasing over time. It extended aquatic lifespan and increased pupal mortality. It also reduced adult lifespan in both sexes with a stronger effect in females. AalDV2 exposure had no significant effect on overall survival, stage-specific mortality, or adult lifespan. However, a significant interaction between viral exposure and thermal stress was detected on aquatic lifespan: AalDV2-exposed females showed further extended larval and pupal development specifically under the 32–34°C regime, without any effect on survival.

These results indicate that the biocontrol potential of AalDV2 cannot be assessed independently of thermal context: while lethal effects were absent under both fluctuating regimes, the prolongation of aquatic development by the virus under thermal stress may have indirect consequences for mosquito population dynamics that warrant further investigation.

**Author Summary:** The tiger mosquito *Aedes albopictus* is among the most invasive species worldwide including in temperate areas and a major vector of dengue, chikungunya, and Zika viruses. Concerns with this species are growing in the context of climate change and effective vector control strategies are increasingly critical. Densoviruses, such as AalDV2, are small DNA viruses that infect mosquitoes and are being explored as potential biocontrol agents because of their restricted host range and ability to spread through shared breeding sites and kill mosquito aquatic stages. However, the understanding of influence of the environmental conditions such as temperature on the effectiveness of these viruses remains incomplete. Here, we investigated the combined effects of fluctuating temperatures (26–28°C and 32–34°C day/night cycles) and AalDV2 exposure on mosquitoes throughout their entire lifespan, from larva to adult death. Our results show that chronic exposure to high fluctuating temperatures strongly reduces mosquito survival, mainly by increasing mortality during pupation and shortening adult lifespan. While AalDV2 exposure did not significantly affect survival or mortality under either temperature regime, a significant interaction between viral exposure and thermal stress was detected: AalDV2-exposed females showed prolonged aquatic development specifically under the high temperature regime (32–34°C). This result demonstrates that temperature shapes the outcome of host–virus interactions even in the absence of lethal effects, and that realistic thermal variability must be incorporated into the evaluation of densovirus-based biocontrol strategies.

## Introduction

Vector-borne diseases pose a major challenge to public health. Aedes-transmitted diseases such as chikungunya, Zika and dengue are re-emerging worldwide and spreading to new regions[1]. Almost half of the global population is exposed to dengue virus with outbreaks multiplying around the world. This geographical expansion coincides with the global spread of *Aedes albopictus* [2], a major vector of several arboviruses and one of the most invasive species worldwide [3]. Originally native to South Asia, *Ae. albopictus* is now established on all continents except Antarctica [4,5]. Its rapid global expansion is largely driven by environmental and climatic changes and facilitated by human activities [6]. The large phenotypic and behavioural plasticity of mosquitoes allows them to fit in urban environments and tolerate wide ranges of temperature [6,7]. On the one hand, anthropogenic landscape modification creates suitable larval habitats, favourable microclimatic conditions, diminishes predation exposure and increases human exposure to vectors. On the other hand, global trade and travel networks allow the introduction of vectors into new regions. As poikilotherms, insects’ distribution is expected to be strongly influenced by climatic conditions. Such thermal fluctuations can alter key life-history traits and consequently modify interactions between *Ae. albopictus*, its pathogens, and human hosts. The life cycle of *Ae. albopictus* is particularly short, therefore repeated exposure to extreme thermal events may substantially affect its survival and population dynamics[8]. Insects are exposed to daily and seasonal thermal fluctuations that can reach stressful levels. To cope with thermal stress, they have evolved many acclimation mechanisms ranging from biomolecular responses such as the expression of Heat Shock Proteins (HSPs) [9,10] to behavioural adjustments in order to maintain their cellular integrity and optimize their survival. Ecological thermal models describe optimal temperature ranges in which moderate increases in temperature are associated with enhanced insect growth and survival, whereas temperatures above or below these optima are detrimental [11]. The optimal temperature range in *Ae. albopictus* is between 21°C and 27°C [12–14]. Larval development is optimal at 29.7°C but slows down at higher temperature, mortality increases beyond 34°C [15,16]. Such sublethal thermal stress may therefore alter physiological trade-offs. One way to mitigate disease transmission is through vector population management therefore several biocontrol tools are being developed to reduce our reliance on insecticide-based methods which are usually non-specific, environmentally toxic and facing extensive resistance mechanisms [17,18].

Concerning mosquito management various approaches are considered : (i) introduction of predators [19] entomopathogenic fungi, bacteria such as *Bacillus thuringiensis* [20] or viruses at all life stages, (ii) the use of olfactory perturbators, (iii) the release of sterile or genetically modified males and use of *Wolbachia*-induced cytoplasmic incompatibility to disrupt reproduction [21–23]. Clearly, because vaccine coverage remains limited and often restricted to specific risk groups, vector control is still the cornerstone of prevention and outbreak response.

Densoviruses (DVs) are single-stranded DNA viruses (Parvoviridae) infecting arthropods and they are considered potential biocontrol agents [22,24]. Their pathogenicity varies depending on both viral strain and host species, although they generally exhibit a relatively restricted host range, generally infecting species within their natural host’s family [24,25]. Infection occurs during the larval stage after which the virus replicates and is released in the water (horizontal transmission)[25] and through transovarian and venereal contact from surviving adults, thereby contaminating oviposition sites. In species displaying skip-oviposition behavior, such as *Ae. albopictus*, DVs may efficiently spread to otherwise inaccessible breeding sites. In a number of lab experiments, AalDV2 has proven to be an efficient biolarvicide for *A. aegypti* but has only moderate effects in *Ae. albopictus* [16]. Additionally, in vitro infection with AalDV2 has been reported to reduce dengue virus (DENV-2) replication, although these results require further investigation [24]. However most mosquito-infecting densoviruses have been discovered and studied in cell cultures and laboratory colonies [24]. Consequently, little is known about the natural distribution of AalDV2, its impact on *Ae. albopictus* biology, the effect of co-infections or the influence of environmental parameters on its virus pathogenicity [24].

In a context of climate change, it is therefore crucial to evaluate how environmental parameters influence densovirus infection dynamics, as these effects may ultimately alter mosquito population size and vectorial capacity. Moreover, the outcome of biological interactions can shift under varying environmental conditions, potentially moving along a continuum from parasitism to mutualism [26]. Such environmentally mediated shifts have been described in plant–virus systems, where viral infection can confer stress tolerance under specific conditions [27–29]. A similar phenomenon could occur in mosquito–densovirus systems. Although densoviruses are primarily studied for their entomopathogenic properties, some have been reported to exhibit mutualistic effects. For example, HaDNV1 infection in the tomato hornworm *Helicoverpa armigera* accelerates development, increases fecundity, and enhances resistance to both baculovirus infection and *Bacillus thuringiensis* toxin used in biocontrol [30].

Previous results showed that AalDV2 infection induces fitness costs, including delayed development and reduced adult body size in *Ae. albopictus*. However, under 34°C thermal stress, exposed individuals exhibited increased survival until adult emergence, suggesting a potential stress tolerance conferred by the AalDV2 infection [16]. As this previous study primarily focused on developmental stages and fixed temperature conditions, we are extending our analysis by measuring survival throughout the entire lifespan under fluctuating temperature. This permits us to get closer to natural conditions associated with daily and seasonal thermal fluctuations as would be the case in a AalDV2-based biocontrol. Specifically, we evaluated the effect of AalDV2 exposure under 32-34°C thermal stress on developmental duration and overall survival until adult death. We hypothesized that AalDV2 exposure enhances thermal tolerance in *Ae. albopictus*, leading to increased survival under heat stress conditions.

## Methods

### Virus production

The AalDV2 strain was purified from a chronically infected cell line of the C6/36 clone[31], which originated from *Ae. albopictus* larvae. This cell line was obtained from the Institut Pasteur in Paris. C6/36 cell lines were cultured at 28°C in RPMI cell culture medium (Gibco, USA), supplemented with 10% heat-inactivated fetal bovine serum (FBS, Gibco, USA), 1% non-essential amino acids (NEAA, Gibco,USA) and 1% penicillin–streptomycin (ATB, Gibco, USA)[25]. The purification started with cell lysis through successive freeze-thaw cycles, then the cells and their supernatants were centrifuged at 4°C for 15 minutes at 3000g to eliminate cell debris. The supernatant was then layered onto a 15% sucrose cushion and subjected to ultracentrifugation at 38 000 rpm (SW41 rotor, Optima^_^90K Beckman Ultracentrifuge) for 3 hours at 8°C. The resulting pellet was resuspended in a 1 mM Tris and 0.1 mM EDTA buffer (TE 0.1X). The final viral suspension was analyzed qualitatively by transmission electron microscopy and quantified by quantitative PCR (qPCR) in genome equivalent virus (gev) per µL as described in Perrin et al (2020)[25]. Control productions were prepared in the same condition, from uninfected C6/36 cells.

### Mosquito maintenance, infection protocol and temperature exposure

We used a strain of *Ae. albopictus* isolated from La Reunion maintained at hundreds of individuals per generation in ISEM, Montpellier, France. The conditions for the experiments were as follows: exposed or not to the virus and, to simulate daily thermal cycles, reared under fluctuation temperatures: a low temperature condition (26°C at night, 28°C during the day) or a high temperature condition (32°C at night, 34°C during the day). Throughout the manuscript, groups are referred to as control/exposed and 26-28°C/32-34°C. Individuals were monitored until their death. These temperatures correspond respectively to optimal and stressful temperatures. We made 2 replicates of the experiment and the temporal delay corresponds to different mosquito generations. Since mortality was expected to be higher at the higher temperature [16] the initial group size was adjusted to ensure sufficient statistical power and representation in each group.

On day 0 (D0) *Ae. albopictus* eggs were submerged in demineralized water to induce eclosion. The following day (D1), hatching larvae were randomly distributed in 96 wells plates, each well containing 180 μL of deionized water.

Half of them (AalDV2-exposed) were exposed to 10^10^ virus equivalent genome (veg) of purified virus (a dose similar to the one used in previous experiments[25] and the other half (Control) to the same volume of an extract of an uninfected C6/36 culture. Plates were distributed in two climatic chambers (T = 26-28°C and 32-34°C, Humidity ~75% and 12:12 light/dark cycle) in which they remained until death. Larvae remained exposed for 24h. On day 3 (D3), larvae were transferred in 12-well plates containing 3 mL of demineralized water in order to ensure sufficient space for larval development. These fluctuating regimes were chosen to approximate warm and hot summer conditions that Aedes albopictus can encounter in container habitats, including during heatwave periods, while remaining within biologically relevant ranges for development and survival. Using fluctuating rather than constant temperatures allowed us to capture thermal variation closer to that experienced in the field and to evaluate whether the fitness effects of AalDV2 observed in constant-temperature assays hold under more realistic conditions.

All mosquitoes had the same food regime of Tetramin baby fish food (0.04 mg/larvae on day 1, 0.06 on day 2, 0.12 on day 3, 0.24 on day 4 and then 0.48mg/larvae every two days). Once pupation happened, pupae were transferred in 10 mL demineralized water in long veiled tubes. After emergence, water was removed from the tubes, through the veil, while the adult was left inside. We discriminated sex based on antenna and proboscis morphology : males have feathery antennae and bifid proboscis [32]. Adult mosquitoes were then fed *ad libitum* on cotton balls soaked in 2% sugar solution for the 4 days following emergence and on cotton balls soaked in demineralized water for the remaining days after the 4th. That food restriction was chosen to narrow energy allocation to the responses against the infection and the heat stress. Dead adults were kept at −20°C, until their infectious state was evaluated. During the experiments, mosquitoes were monitored every day and date was recorded for every event (death, pupation and emergence).

### Virus detection in mosquitoes

The qPCR was performed using the LightCycler 480 System software (Roche, France) for 45 cycles (5 sec denaturation 95°C, 10 sec hybridization 62°C) and the SensiFAST SYBR No-ROX Kit (Meridian Biosciences). All DNA samples were obtained via phenol chloroform extraction and they were analyzed in triplicates for each quantification. For AalDV2 and the actin the following protocol was used: 5 min at 95°C, 30 cycles of 94°C for 30 s, 52°C for 30 s, and 72°C for 60 s, then 72°C for 5 min. The primers for the amplification of the *Ae. albopictus’* actine primers were actAlb-dir: 5’-GCAAACGTGGTATCCTGAC-3’and actAlb-rev: 5’-GTCAGGAGAACTGGGTGCT-3’ [43]. For AalDV2 the primers were qAalDV2_AP_2_F: 5’-CTCTGGAGCCGCTGTGTAAT-3’ and qAalDV2_AP_2_R:5’ TGGCCAACAATTACGAACAA-3’ [36]. Following a DNA-based relative quantification approach, the ratio between the AalDV2 and actin Ct values was used to estimate the relative number of viral genome copies per mosquito genome equivalent. To rule out potential cross-contamination, 10 mosquitoes were randomly selected in each non-exposed groups and tested for viral infection. All unexposed controls tested negative and were then considered non-infected. Regarding the exposed mosquitoes only a fraction of them were tested for the presence of the virus as a quality control of the infection protocol. If the presence of the virus reflects an infection, its absence after exposure in the adult does not mean that the infection has not occurred as it may have been cleared by the mosquito immune system. Accordingly, we’ll use the term exposed vs unexposed in the manuscript.

### Statistical analysis

Statistical analyses were performed in R (v4.5.0) using the following packages: survival, lme4, glmmTMB, emmeans, DHARMa, FSA, and performance. A significance threshold of 0.05 was applied throughout. When permitted by the experimental design, the replicate was used as a random effect. Model assumptions were assessed using standard diagnostic procedures and pairwise comparisons were conducted using estimated marginal means (emmeans) with Šidák adjustment for multiple comparisons.

Overall survival was analysed using Cox proportional hazards models including temperature, treatment, and their interaction as predictors. Violation of the proportional hazards assumption for temperature was addressed by including a time-dependent effect [44]. Global model significance was assessed via likelihood ratio test, and predictive performance was assessed through the concordance index (C-index).

Stage-specific mortality (larval, pupal, adult) was analysed using binomial generalized linear mixed models (GLMMs) with treatment and temperature as fixed effects.

Aquatic lifespan was analysed using GLMMs with a Gamma distribution and log link function, including temperature, treatment, sex, and their interactions. Minor deviations from normality were considered acceptable given that overdispersion, zero-inflation, and dispersion tests were all satisfactory. Results were confirmed with a non-parametric Kruskal-Wallis test followed by Dunn’s post-hoc test with Bonferroni correction.

Adult lifespan was analysed using linear mixed models after square-root transformation of the response variable, with temperature, treatment, sex, and their interactions as fixed effect. Minor departures from normality were acceptable given all groups exceeded 30 individuals, providing robustness to normality violations and heteroscedasticity remained acceptable.

## Results

### Effect of temperature on survival and development

Exposure to 32–34°C significantly increased mortality risk and reduced median lifespan by approximately 10 days (Fig. 1). This effect intensified over time, as indicated by a significant temperature × time interaction (HR = 2.21, p = 0.0018). The proportional hazards assumption was violated for temperature (Schoenfeld test: p = 0.0024). This confirms a time-dependent effect in which the detrimental impact of chronic exposure to 32-34°C intensifies (Fig. 1). Adult lifespan was markedly reduced at 32–34°C (χ^2^ = 66.29, df = 1, p < 0.001), particularly in females (Fig. 3). Mean adult lifespan ranged from approximately 20–25 days at 26-28°C to 7–15 days at 32-34°C (Fig. 3).

**Fig 1:**
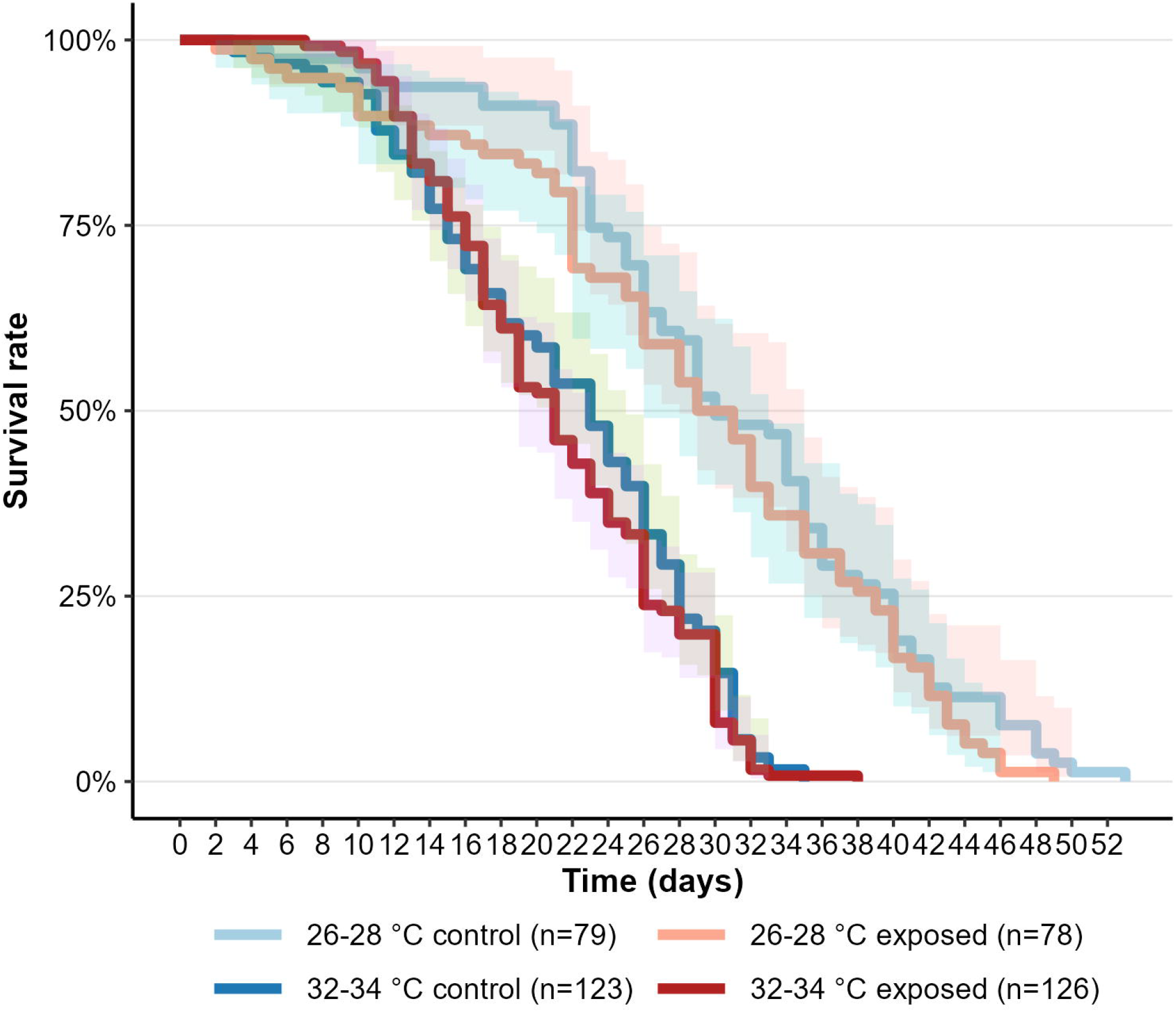
Overall survival curves of *Aedes albopictus* adults according to temperature regime (26–28°C vs. 32–34°C) and AalDV2 exposure treatment (control vs. exposed). Survival was monitored daily from emergence until death. Shaded areas represent 95% confidence intervals. Total sample sizes: 26–28°C control n = 79, 26–28°C exposed n = 78, 32–34°C control n = 123, 32–34°C exposed n = 126.

In contrast, aquatic lifespan increased by approximatively 2.5 days (Fig. 4), reflecting an extended larval stage, while pupal duration remained constant. This prolongation of larval development was associated with increased pupal mortality, rising significantly (p = 0.001) from 2.5% at 26–28°C to 13.8% at 32–34°C (Fig. 2). Individuals that failed to complete development survived on average between 10 and 13 days, with divergence in survival probabilities becoming apparent around day 13 post-hatching (Fig. 1). Emergence consistently occurred between days 9 and 16 across all groups.

**Fig 2:**
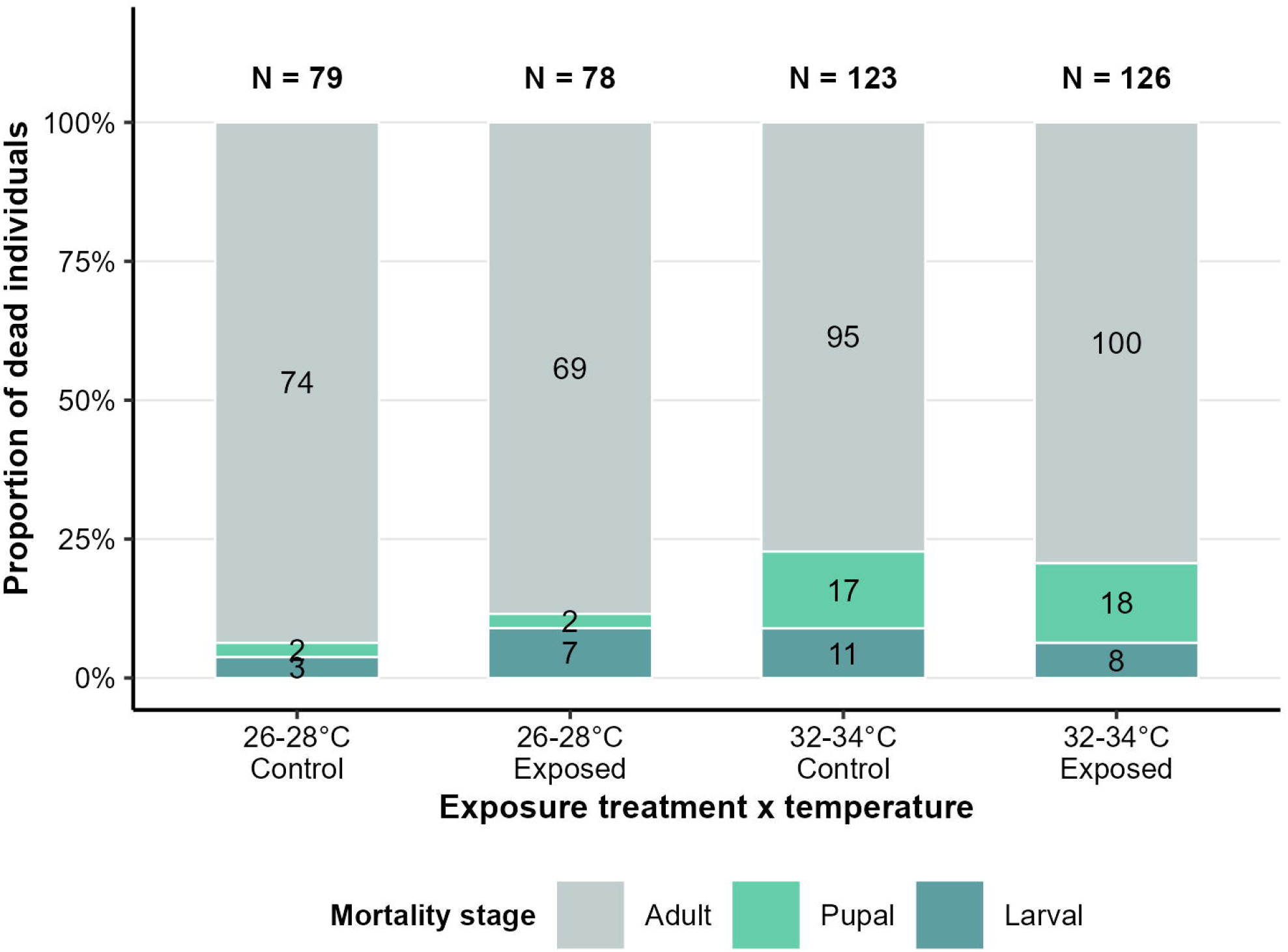
Proportion of dead individuals at each developmental stage (larval, pupal, adult) according to temperature regime and AalDV2 exposure treatment. Numbers above bars indicate the total number of dead individuals per group. Total individuals monitored: 26–28°C control n = 79, 26–28°C exposed n = 78, 32–34°C control n = 123, 32–34°C exposed n = 126. Note the marked increase in pupal mortality under the high temperature regime (32–34°C), regardless of exposure treatment

**Fig 3:**
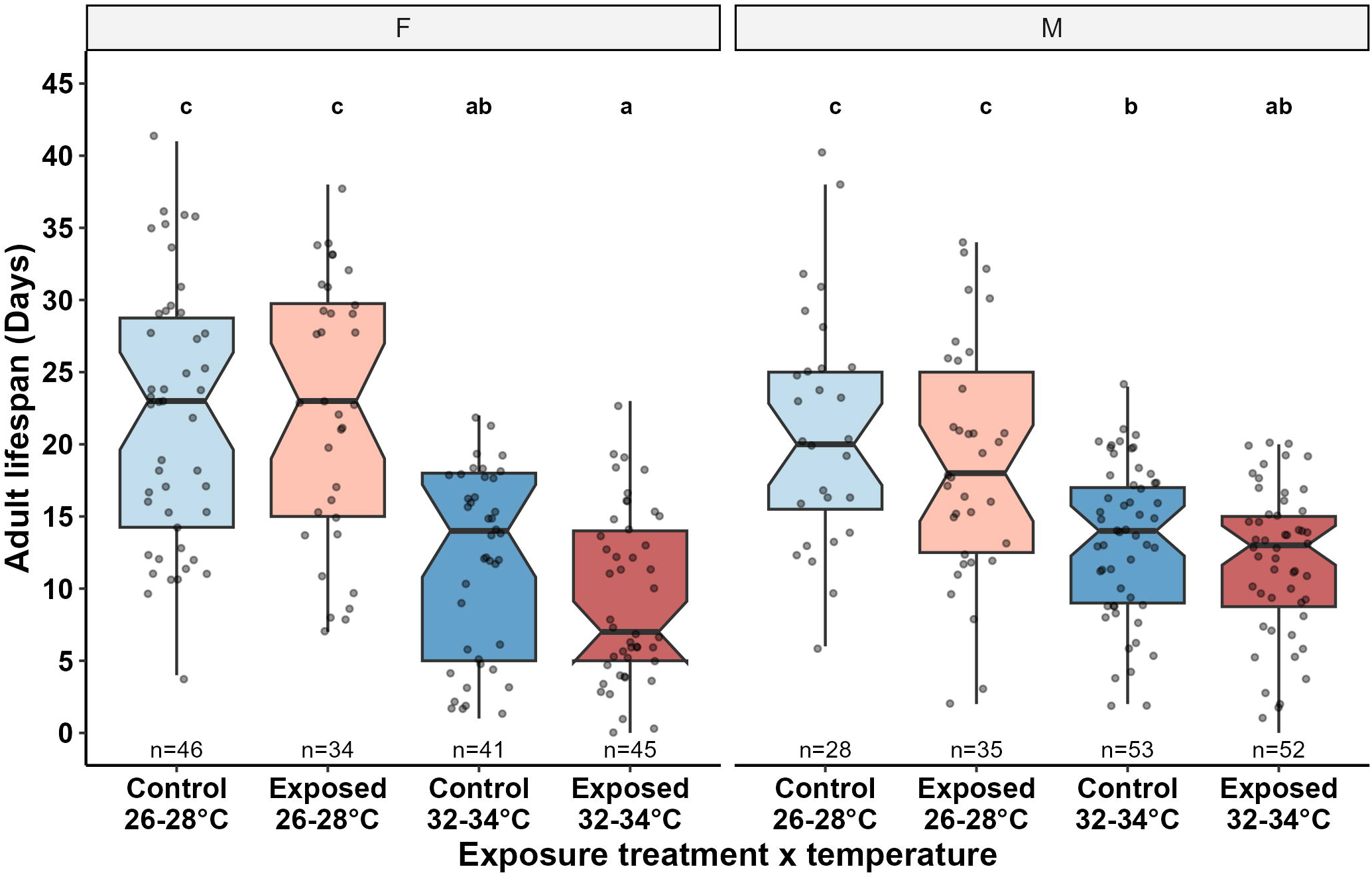
Adult lifespan (days) of *Aedes albopictus* females (F) and males (M) according to temperature regime and AalDV2 exposure treatment. Boxplots show median (centre line), interquartile range (box), and 1.5× IQR (whiskers); individual data points are overlaid. Letters above boxplots indicate results of pairwise post-hoc comparisons (Šidák correction); groups sharing a letter do not differ significantly (p > 0.05). Sample sizes per group are indicated above each boxplot. Females: 26–28°C control n = 46, 26–28°C exposed n = 34, 32–34°C control n = 41, 32–34°C exposed n = 45. Males: 26–28°C control n = 28, 26–28°C exposed n = 35, 32–34°C control n = 53, 32–34°C exposed n = 52

**Fig 4:**
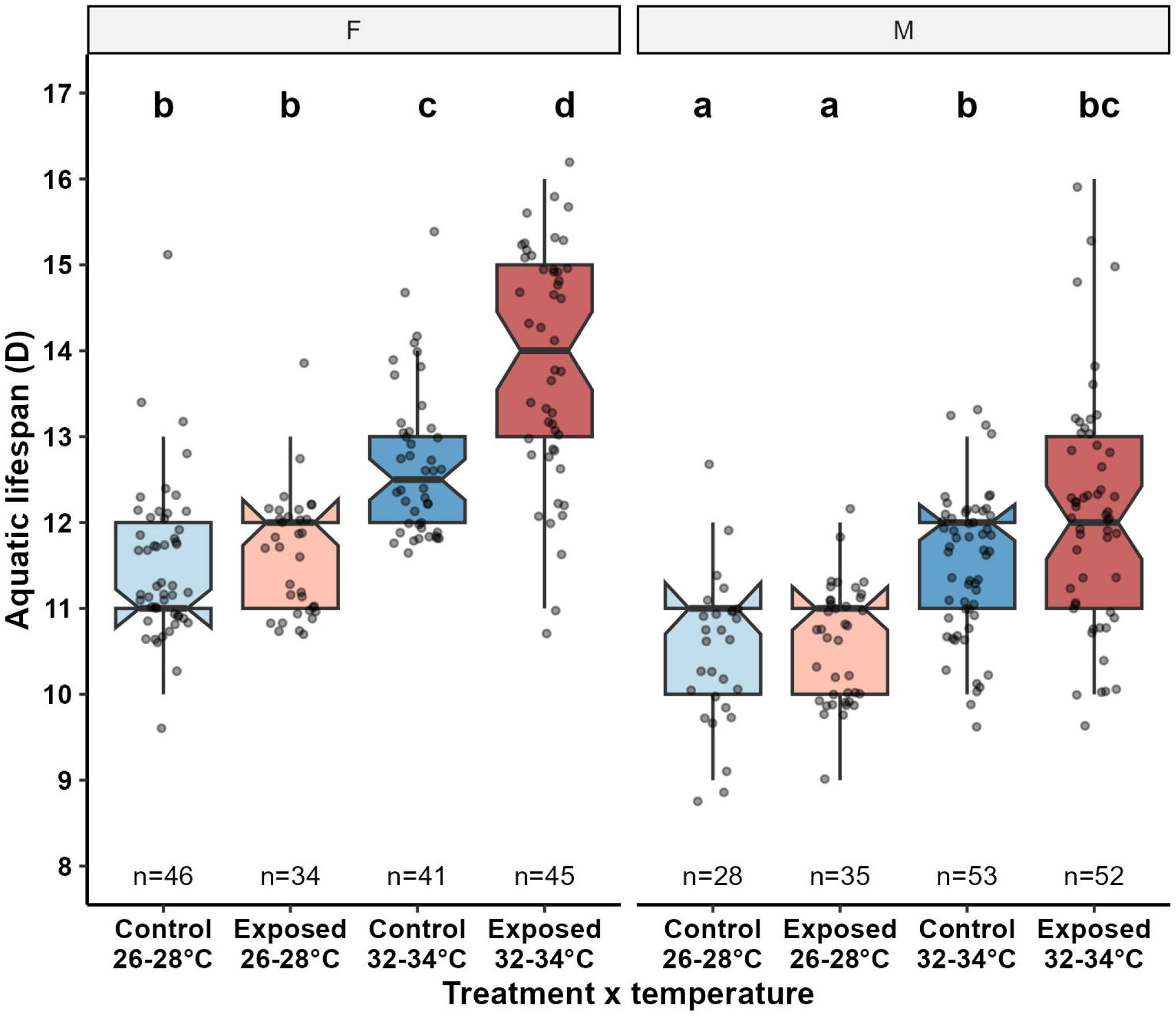
Aquatic lifespan (days) of *Aedes albopictus* females (F) and males (M) according to temperature regime and AalDV2 exposure treatment. Boxplots show median (centre line), interquartile range (box), and 1.5× IQR (whiskers); individual data points are overlaid. Letters above boxplots indicate results of pairwise post-hoc comparisons (Šidák correction); groups sharing a letter do not differ significantly (p > 0.05). Sample sizes per group are indicated above each boxplot. Females: 26–28°C control n = 46, 26–28°C exposed n = 34, 32–34°C control n = 41, 32–34°C exposed n = 45. Males: 26–28°C control n = 28, 26–28°C exposed n = 35, 32–34°C control n = 53, 32–34°C exposed n = 52.

### Effect of AalDV2 exposure on survival and development

AalDV2 exposure had no significant effect on overall survival (HR = 1.21, p = 0.232), stage-specific mortality, or adult lifespan (all p > 0.2).

### Combined effect of AalDV2 exposure and chronic thermal stress on survival and development of Ae. albopictus

While AalDV2 exposure did not affect survival traits (see above) when combined with chronic thermal stress it increased aquatic lifespan (z = 2.47, p = 0.013), indicating that the effect of exposure on developmental duration depended on thermal conditions. Post-hoc comparisons revealed that exposed individuals at 32-34°C exhibited increased aquatic duration relative to controls. No interactions involving sex were detected (all p > 0.30).

Larval survival remained high (>90%) and was not affected by temperature, viral exposure, or their interaction (all p > 0.1; Fig. 2). However, females exposed to both AalDV2 and 32-34°C exhibited a significantly longer aquatic lifespan compared to control females under the same thermal condition (Fig. 4).

#### Sex ratio

Temperature had a significant effect on sex ratio (p = 0.0176), with a shift from female-biased at 26–28°C (0.6) to male-biased at 32–34°C (1.3) (Fig. 5), although effect sizes were small and pairwise comparisons were not significant. AalDV2 exposure had no significant effect on sex ratio, nor did its interaction with temperature.

**Fig 5:**
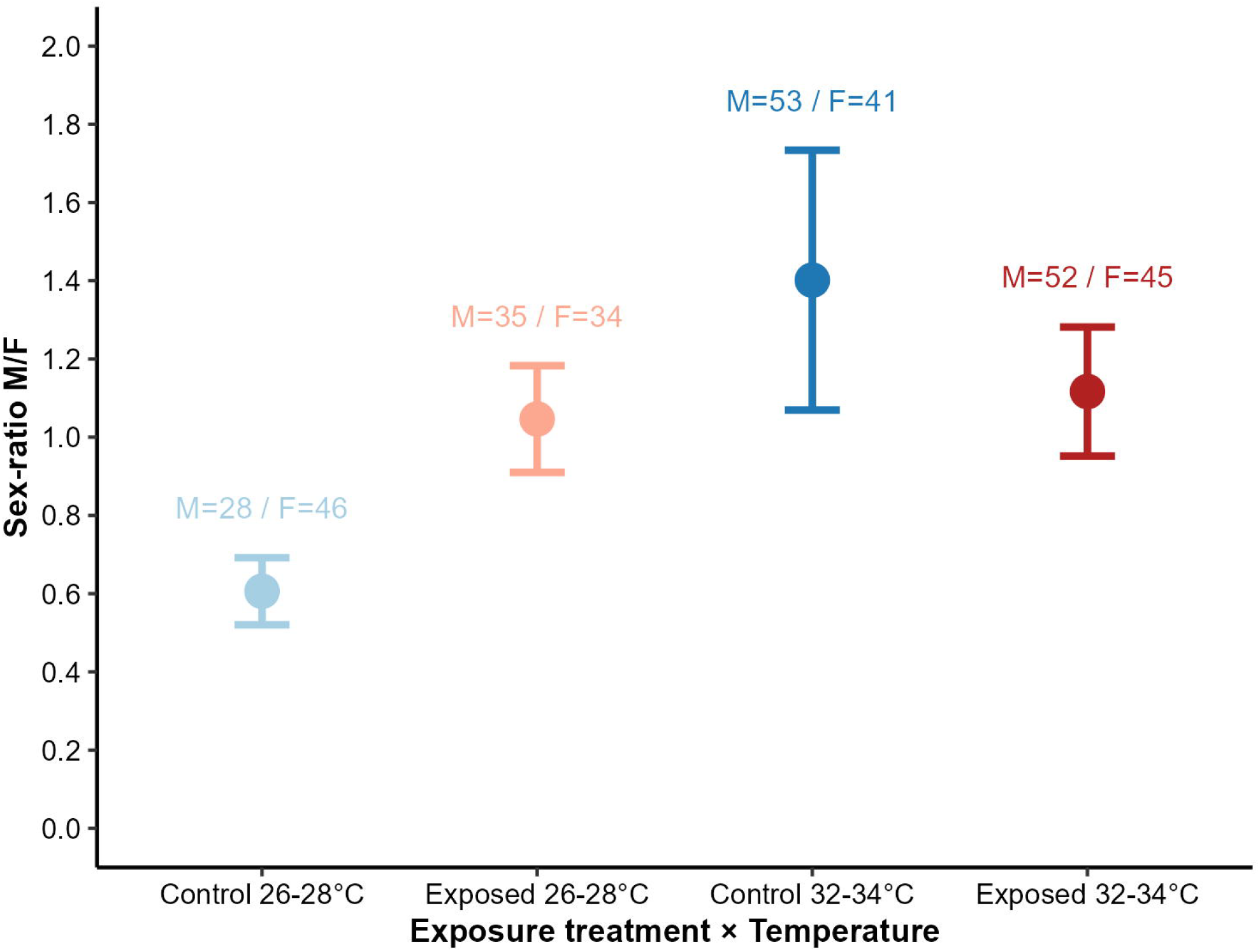
Sex ratio (males/females) of emerging *Aedes albopictus* adults according to temperature regime and AalDV2 exposure treatment. Each point represents the sex ratio observed in one replicate tube; error bars indicate ± standard deviation across replicates. Letters above groups indicate results of pairwise post-hoc comparisons; groups sharing a letter do not differ significantly (p > 0.05). Numbers of males and females emerging per group are indicated within or above each bar. Total individuals: 26–28°C control n = 74, 26–28°C exposed n = 69, 32–34°C control n = 94, 32–34°C exposed n = 97. A sex ratio of 1.0 (dashed line) indicates equal proportions of males and females

Sex also significantly influenced aquatic lifespan (z = −4.76, p < 0.001) with females having a longer aquatic development than males across conditions. Regarding adult lifespan, the interaction between temperature and sex showed a marginal trend (χ^2^ = 3.03, df = 1, p = 0.082), with a reduction in lifespan at high temperature more pronounced in females than in males. (Fig 4). However post-hoc pairwise comparisons (Šidák correction) confirmed that temperature was the primary driver of differences among groups.

## Discussion

Our study evaluated the combined effect of temperature and AalDV2 exposure on the survival and development across the complete life cycle of *Ae. albopictus*. Our results indicate that chronic thermal stress strongly impaired survival and development, whereas AalDV2 exposure and its interaction with temperature has limited effects under the tested conditions. These findings highlight that under warm and hot fluctuating conditions, temperature is the primary driver of mosquito performance, and that the fitness costs of AalDV2 exposure are negligible, at least for the life-history traits measured here.

### Chronic thermal stress

The reduction in overall survival under 32-34°C likely resulted from a dual effect of temperature: a higher mortality during pupation and a reduced adult longevity. Thermal stress may disrupt the pupation process itself, a complex, energy-intensive process regulated by the steroid hormone ecdysone in response to physiological cues (larval size, nutritional status) and external signals (photoperiod) [34]. Previous work in *Drosophila melanogaster* has shown that heat stress reduces larval weight and decreases the probability of successful pupation[35]. Chronic exposure to 32-34°C may also induce heat shock proteins (HSPs) expression[10], proteins involved in mechanisms of tolerance to high thermal stress [36]. These adaptive responses have a significant metabolic cost that could limit resources normally allocated to pupation. Furthermore, HSP synthesis is coupled to the ecdysteroid peak under standard conditions, and thermal stress is likely to disrupt this co-regulation [37], consequently compromising metamorphosis. Monitoring larval development by measuring metabolic parameters (e.g., larval weight) and relevant protein parameters in response to chronic stress would help shed light on these mechanisms directly.

### AalDV2 exposure and combined effect with temperature

AalDV2 exposure alone did not significantly affect overall survival stage-specific mortality, or adult lifespan (Fig 2-D). This contrasts with a recent study conducted under constant and high temperature 34°C [16]. In this latter one AalDV2 exposure conferred thermal tolerance and an improved survival to adult emergence. That protective effect is absent in the current study under fluctuating temperatures. This difference is unlikely to be attributed to differences in viral strain or mosquito colony, as we used the same biological material. Nevertheless, we cannot exclude changes over time in the mosquito colony of the viral stocks (e.g., laboratory adaptation, genetic drift, or variation in passage history), which could affect infection dynamics and virulence.

Despite the absence of effects on survival, a significant interaction temperature × AalDV2 exposure interaction was detected for aquatic lifespan (z = 2.47, p = 0.013). Interestingly at 32–34°C females exhibited a prolonged aquatic development relative to unexposed controls. This developmental delay was not accompanied by increased larval or pupal mortality, suggesting a sublethal effect of the virus–temperature interaction on development rather than on survival per se.

From a vector control perspective, the absence of strong fitness costs under fluctuating temperature regimes suggests that AalDV2 alone may have limited capacity to suppress *Ae. albopictus* populations in warm environments. This result highlights the fact that densovirus-based interventions should not be evaluated solely under constant laboratory conditions and that their contribution to integrated vector management is likely to be context dependent on local thermal regimes.

### Sex specific responses

Thermal stress appeared to differentially affect male and female developmental trajectories. This shift in sex ratio from female biased to male-biased remains modest but it could reflect sex-specific differences in physiological sensitivity during late larval or pupal stages. Given that female aquatic lifespan is generally longer than that of males [49], development under thermal stress may disproportionately increase female mortality before emergence, thereby reducing female proportion at emergence.

Females exposed to both viral infection and high temperature exhibited extended aquatic lifespan without increased mortality (Fig. 4, 5). This suggests that females may be more sensitive to environmental stressors, possibly due to differences in energy allocation and reproductive physiology. Future studies using molecular sexing approaches (e.g., Fru gene detection) during larval stages would help clarify the timing and mechanisms underlying these sex-specific responses.

Although we did not directly measure arbovirus infection or transmission in this study, our results have implications for *Aedes*-borne disease risk. Survival, development time and adult lifespan are key components of vectorial capacity, and our data indicate that chronic heat stress at 32–34°C can substantially reduce these traits, potentially limiting transmission in very hot conditions. At the same time, the negligible additional impact of AalDV2 on these traits under fluctuating temperatures suggests that densovirus-based biocontrol may contribute little to reducing vectorial capacity in warm regions. As climate change increases the frequency and intensity of heatwaves in many areas where *Aedes* albopictus is established, incorporating realistic thermal variation into both experimental assays and transmission models will be essential to accurately predict the performance of viral biocontrol tools and their potential role in strategies targeting dengue, chikungunya and Zika.

Contrary to our initial hypothesis, AalDV2 infection did not confer any measurable thermal tolerance advantage under the fluctuating temperature regimes. While our findings differ from those previously reported [25,36], they provide complementary insights into the plasticity of *Ae. albopictus* responses to environmental stressors. Unlike an earlier study, conducted under fixed temperatures (28°C and 34°C) this work used fluctuating thermal regimes (26–28°C and 32–34°C, day/night cycles). This approach revealed an unexpected resilience in mosquitoes under fluctuating thermal stress conditions with lower mortality rates (below 25 %, Fig. 1) compared to up to 70% under constant 34°C[16]. This is consistent with findings in other insects. For instance, in *Lycaena tityrus*, fluctuating temperature led to shorter development times, increased heat stress resistance, decreased HSPs expression, and increased immunocompetence compared to constant temperatures [39]. Therefore, the difference in our experimental design may substantially influence our result.

As with any controlled laboratory study or modelling approach, several limitations need to be acknowledged. Thus, the experiment used a single viral dose and a single colony established from La Réunion island, reflecting a deliberate choice to work under controlled and reproducible conditions. Generalization to other AalDV2 strains, doses, or mosquito populations with distinct thermal histories are then limited. The relatively small group sizes when subdivided by sex and condition (n ≤ 60) may have limited statistical power to detect minor effects of AalDV2 exposure.

Adult mosquitoes were maintained on a sugar solution for several days then on water only, without blood meals. Clearly this limits the relevance of adult survival data and precludes any assessment of fecundity or vectorial capacity that are key parameters for evaluating the efficacy of a biocontrol agent. Finally, our experiment had temporal replication with two experimental blocks but using a fully independent colony would strengthen the generalizability of our conclusions.

Knowing that most mosquito-infecting densovirus have been discovered and studied in cell cultures and laboratory colonies[24] more experiments should be considered before using them in natural conditions. Furthermore, in both our experiments, AalDV2 did not affect mortality under control conditions indicating its potential use as a biocontrol agent for *Ae. albopictus* might not be as efficient as expected. We cannot conclude whether or not the effect of the interaction between temperature and infection on the aquatic lifespan might be positive for population dynamics, therefore models including data on the consequences of AalDV2 infection on such life history traits.

## Conclusion

Overall, our findings have important implications for a better understanding of mosquito-virus interactions but also in the more applied field of vector control. They suggest that the performance of AalDV2 cannot be evaluated independently of environmental context and point out the importance of fluctuating thermal regimes that may substantially alter the outcome of host–virus interactions compared to constant laboratory conditions. By revealing the absence of impact on mosquito mortality, our work raises questions about the efficacy of AalDV2 as a biocontrol tool for *Ae. albopictus*. In a more general view, most mosquito-infecting densoviruses have been discovered and studied in cell cultures and laboratory colonies[24] and field-relevant evaluation under realistic thermal variability are a necessary step before considering their eventual use them in natural conditions. The incorporation of the sublethal effects of AalDV2 infection on life-history traits of mosquito vectors in modelling approach would also be a crucial point to consider. This would help determining how they could translate into real impacts on mosquito population and arbovirus transmission, a critical point for public health.

## Ethical statement

This study involved standard laboratory procedures with mosquito larvae and cell cultures. No specific ethical approval is required under French institutional regulations for invertebrate laboratory research. All procedures were conducted in compliance with the institutional biosafety guidelines of ISEM, Université de Montpellier.

## Author contributions

Conceptualization: N.S. C.B. Data curation: N.S., C.B. Formal analysis: N.S., C.B. Funding acquisition: C.B, MVM. Investigation: N.S., M.P.-S., P.M. Methodology: N.S., M.P.-S., P.M., A-S G.G., M.V.M., A.M., C.B. Project administration: C.B. Supervision: C.B. Visualization: N.S., C.B. Writing – original draft: N.S. Writing – review & editing: C.B.

## Acknowledgements

We are grateful to Doriane Mutuel for the maintenance of the cell cultures and to Anne-Sophie Gosselin-Grenet for the AalDV2 production. This work was supported by the French ANSES Environment-Santé-Travail research program (PNR EST) (project 2018/1/183, “DENSOTOOL”, 2019–2023 to CB) and by the initiative CLAPAS (IdeTT 2025-2026 to CB, MVM and NS)

## Data availability

## Figures

**Table 1:** Cox proportional hazards model for overall survival including temperature (32–34°C vs. 26–28°C), AalDV2 exposure, their interaction, and a time-dependent term for temperature to account for the violation of the proportional hazards assumption (Schoenfeld test: p = 0.0024). Hazard ratios (HR) > 1 indicate increased mortality risk. n= 406 events, model concordance index = 0.631, likelihood ratio test χ^2^=123.9, df=4, p<2×10-^16^. Significance codes: ** p < 0.01.

| Effect | Hazard Ratio (HR) | 95% CI | Pr(>\|z\|) |
| --- | --- | --- | --- |
| Temperature (32-34°C) | 0.33 | 0.07-1.53 | 0.155 |
| AalDV2 exposure | 1.21 | 0.88-1.67 | 0.232 |
| Temperature x AalDV2 exposure | 0.92 | 0.62-1.39 | 0.0705 |
| Temperature (32-34°C) x<br>log(time) | 2.21 | 1.35-3.63 | 0.00175 ** |

**Table 2:**
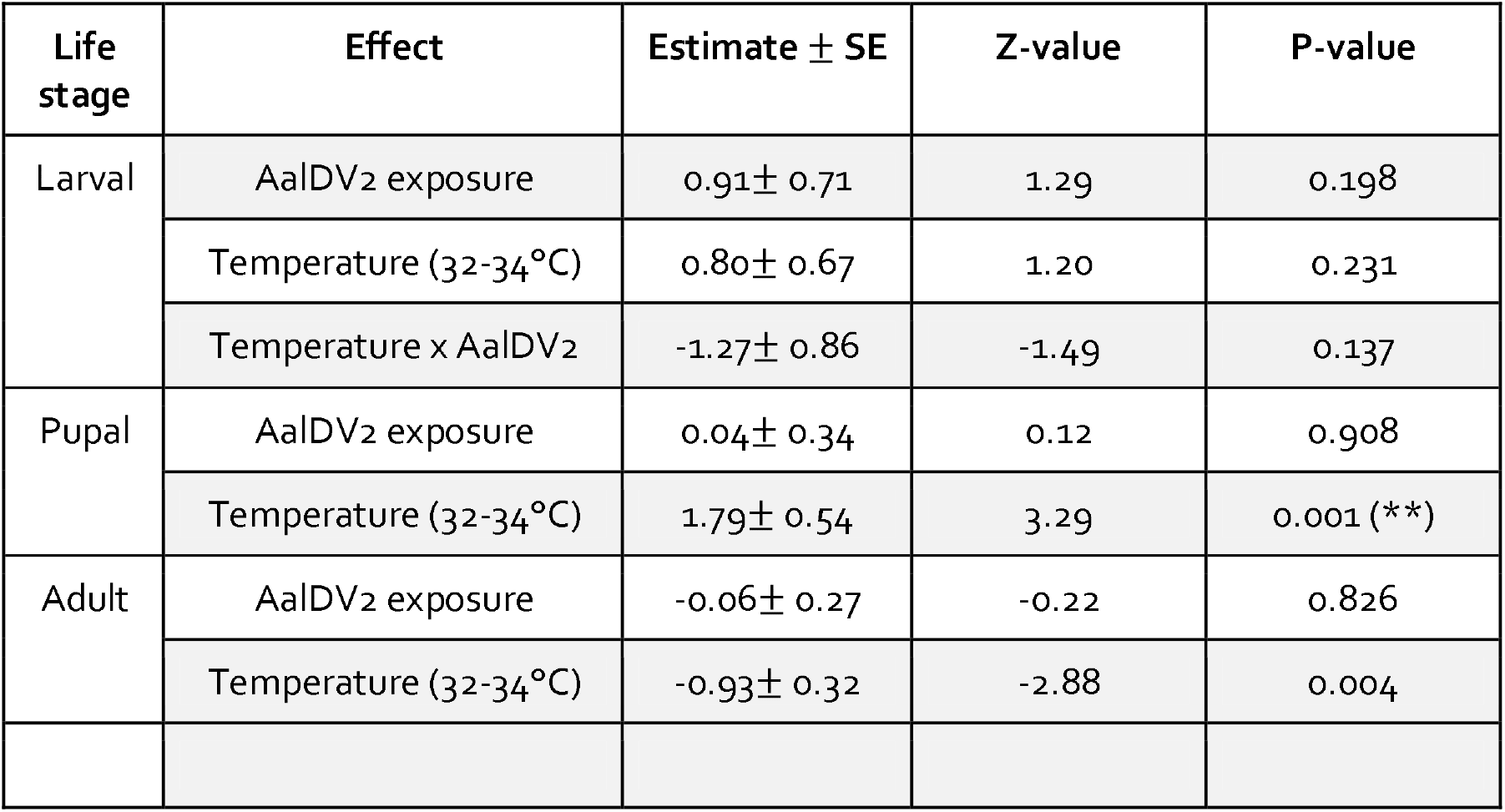
Binomial Generalized Linear Mixed Models (GLMMs) for stage specific mortality larval, pupal, and adult). Replicate was used as a random intercept. The temperature × AalDV2 interaction term was included for larval mortality only; it was not estimated for pupal and adult stages due to insufficient variation in the response variable. Significance codes: ** p < 0.01.

**Table 3:**
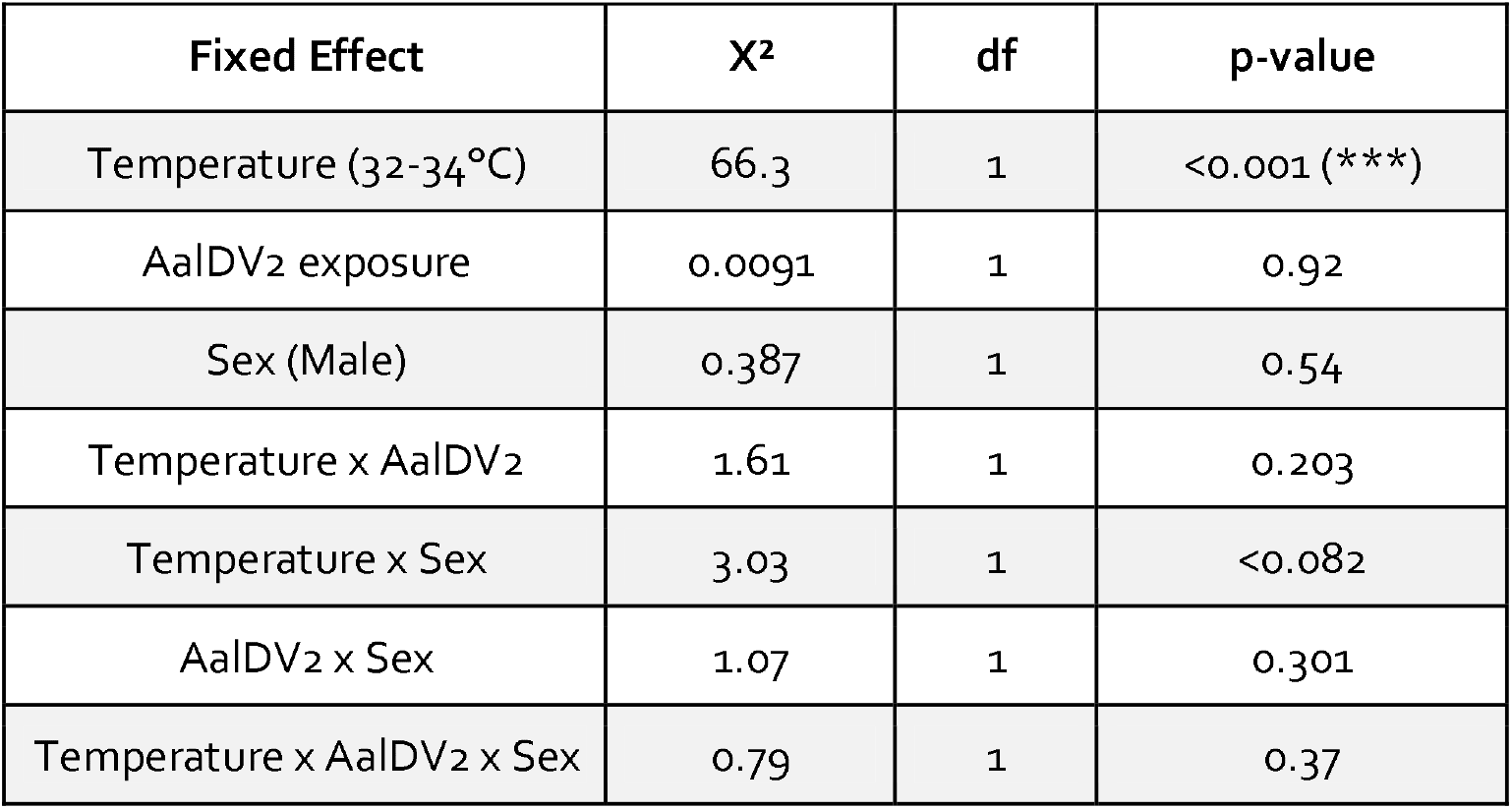
Linear mixed-effects model (LMM) for adult lifespan testing the effects of temperature, viral exposure (AalDV2), sex, and their interactions. Replicate was included as a random intercept. Wald χ^2^ tests correspond to Type III sums of squares. Significance codes: *** p < 0.001.

| Fixed Effect | X<sup>2</sup> | df | p-value |
| --- | --- | --- | --- |
| Temperature (32-34°C) | 66.3 | 1 | <0.001 (***) |
| AalDV2 exposure | 0.0091 | 1 | 0.92 |
| Sex (Male) | 0.387 | 1 | 0.54 |
| Temperature x AalDV2 | 1.61 | 1 | 0.203 |
| Temperature x Sex | 3.03 | 1 | <0.082 |
| AalDV2 x Sex | 1.07 | 1 | 0.301 |
| Temperature x AalDV2 x Sex | 0.79 | 1 | 0.37 |

**Table 4:** Generalized linear mixed model (Gamma distribution, log link) for aquatic lifespan testing the effects of temperature, viral exposure (AalDV2), sex, and their interactions. Replicate was included as a random intercept. Post-hoc pairwise comparisons were performed using the Šidák adjustment method (emmeans package, α = 0.05). Results were confirmed using non-parametric Kruskal-Wallis testing (p < 2.2e-16) followed by Dunn’s test with Bonferroni correction.

| Fixed effect | Estimate ± SE | Z-Value | P-value |
| --- | --- | --- | --- |
| Temperature (32-34°C) | 0.091 ± 0.017 | 5.23 | <0.001 (***) |
| AalDV2 exposure | 0.022 ± 0.018 | 1.19 | 0.233 |
| Sex (male) | -0.092 ± 0.019 | -4.76 | <0.001 (***) |
| Temperature x AalDV2 | 0.062 ± 0.025 | 2.47 | 0.013 (*) |
| Temperature x Sex | 0.006 ± 0.0026 | -0.23 | 0.817 |
| AalDV2 x Sex | -0.019 ± 0.027 | -0.70 | 0.486 |
| Temperature x Sex x AalDV2 | 0.017 ± 0.036 | -0.47 | 0.640 |

**Table 5:** Binomial GLMM for sex ratio. Although temperature showed a statistically significant effect, pairwise comparisons with Bonferroni correction revealed no significant differences. This result was confirmed by a Kruskal wallis test (χ^2^ = 5.50, df=3, p=0.139).

| Fixed Effect | Estimate SE | Z-value | p-value |
| --- | --- | --- | --- |
| AalDV2 exposure | 0.53 0.34 | 1.55 | 0.122 |
| Temperature (32-34°C) | 0.75 0.32 | 2.37 | 0.018 (*) |
| AalDV2 exposure x Temperature | -0.64 0.45 | -1.43 | 0.154 |

## Bibliography

1. McKenzie BA, Wilson AE, Zohdy S. *Aedes albopictus* is a competent vector of Zika virus: A meta-analysis. PLOS ONE. 2019;14: e0216794. doi:10.1371/journal.pone.0216794

2. Skuse FAA. The banded mosquito of Bengal. Indian Mus Notes. 1895.

3. Bonizzoni M, Gasperi G, Chen X, James AA. The invasive mosquito species *Aedes albopictus*: current knowledge and future perspectives. Trends Parasitol. 2013;29: 460–468. doi:10.1016/j.pt.2013.07.003

4. Goubert C, Minard G, Vieira C, Boulesteix M. Population genetics of the Asian tiger mosquito *Aedes albopictus*, an invasive vector of human diseases. Heredity. 2016;117: 125–134. doi:10.1038/hdy.2016.35

5. Kraemer MU, Sinka ME, Duda KA, Mylne AQ, Shearer FM, Barker CM, et al. The global distribution of the arbovirus vectors *Aedes aegypti* and *Ae. albopictu*s. Jit M, editor. eLife. 2015;4: e08347. doi:10.7554/eLife.08347

6. Barman S, Semenza JC, Singh P, Sjödin H, Rocklöv J, Wallin J. A climate and population dependent diffusion model forecasts the spread of *Aedes albopictus* mosquitoes in Europe. Commun Earth Environ. 2025;6: 1–12. doi:10.1038/s43247-025-02199-z

7. Couper LI, Dodge TO, Hemker JA, Kim BY, Exposito-Alonso M, Brem RB, et al. Evolutionary adaptation under climate change: *Aedes* sp. demonstrates potential to adapt to warming. Proc Natl Acad Sci. 2025;122: e2418199122. doi:10.1073/pnas.2418199122

8. de Souza WM, Weaver SC. Effects of climate change and human activities on vector-borne diseases. Nat Rev Microbiol. 2024;22: 476–491. doi:10.1038/s41579-024-01026-0

9. King AM, MacRae TH. Insect Heat Shock Proteins During Stress and Diapause. Annu Rev Entomol. 2015;60: 59–75. doi:10.1146/annurev-ento-011613-162107

10. Sivan A, Shriram AN, Muruganandam N, Thamizhmani R. Expression of heat shock proteins (HSPs) in *Aedes aegypti* (L) and *Aedes albopictus* (Skuse) (Diptera: Culicidae) larvae in response to thermal stress. Acta Trop. 2017;167: 121–127. doi:10.1016/j.actatropica.2016.12.017

11. Damos P, Savopoulou-Soultani M. Temperature-Driven Models for Insect Development and Vital Thermal Requirements. Psyche J Entomol. 2012;2012: 123405. doi:10.1155/2012/123405

12. Schmidt CA, Comeau G, Monaghan AJ, Williamson DJ, Ernst KC. Effects of desiccation stress on adult female longevity in *Aedes aegypti* and *Ae. albopictus* (Diptera: Culicidae): results of a systematic review and pooled survival analysis. Parasit Vectors. 2018;11: 267. doi:10.1186/s13071-018-2808-6

13. Alto BW, Juliano SA. Temperature Effects on the Dynamics of *Aedes albopictus* (Diptera: Culicidae) Populations in the Laboratory. J Med Entomol. 2001;38: 548–556. doi:10.1603/0022-2585-38.4.548

14. Marini G, Manica M, Arnoldi D, Inama E, Rosà R, Rizzoli A. Influence of Temperature on the Life-Cycle Dynamics of *Aedes albopictus* Population Established at Temperate Latitudes: A Laboratory Experiment. Insects. 2020;11: 808. doi:10.3390/insects11110808

15. Reinhold JM, Lazzari CR, Lahondère C. Effects of the Environmental Temperature on *Aedes aegypti* and *Aedes albopictus* Mosquitoes: A Review. Insects. 2018;9: 158. doi:10.3390/insects9040158

16. Boëte C, Perriat-Sanguinet M, Gosselin-Grenet A-S, Makoundou P, Ogliastro M, Sicard M, et al. Stranger Swings: Temperature-Dependent Upsides and Downsides of a Densovirus in *Aedes albopictus*. bioRxiv; 2026. p. 2026.03.09.710552. doi:10.64898/2026.03.09.710552

17. Zulfa R, Lo W-C, Cheng P-C, Martini M, Chuang T-W. Updating the Insecticide Resistance Status of *Aedes aegypti* and *Aedes albopictus* in Asia: A Systematic Review and Meta-Analysis. Trop Med Infect Dis. 2022;7. doi:10.3390/tropicalmed7100306

18. Vontas J, Kioulos E, Pavlidi N, Morou E, della Torre A, Ranson H. Insecticide resistance in the major dengue vectors *Aedes albopictus* and *Aedes aegypti*. Pestic Biochem Physiol. 2012;104: 126–131. doi:10.1016/j.pestbp.2012.05.008

19. Udayanga L, Ranathunge T, Iqbal MCM, Abeyewickreme W, Hapugoda M. Predatory efficacy of five locally available copepods on *Aedes* larvae under laboratory settings: An approach towards bio-control of dengue in Sri Lanka. PLOS ONE. 2019;14: e0216140. doi:10.1371/journal.pone.0216140

20. Land M, Bundschuh M, Hopkins RJ, Poulin B, McKie BG. Effects of mosquito control using the microbial agent *Bacillus thuringiensis israelensis* (Bti) on aquatic and terrestrial ecosystems: a systematic review. Environ Evid. 2023;12: 26. doi:10.1186/s13750-023-00319-w

21. Benelli G, Jeffries CL, Walker T. Biological Control of Mosquito Vectors: Past, Present, and Future. Insects. 2016;7: 52. doi:10.3390/insects7040052

22. Abd-Alla AMM, Meki IK, Demirbas-Uzel G. Insect Viruses as Biocontrol Agents: Challenges and Opportunities. In: El-Wakeil N, Saleh M, Abu-hashim M, editors. Cottage Industry of Biocontrol Agents and Their Applications: Practical Aspects to Deal Biologically with Pests and Stresses Facing Strategic Crops. Cham: Springer International Publishing; 2020. pp. 277–295. doi:10.1007/978-3-030-33161-0_9

23. Fauvergue X, Rusch A, Barret M, Bardin M, Jacquin-Joly E, Malausa T, et al. Biocontrôle. Editions Quae; 2020. Available: https://hal.inrae.fr/hal-02791036

24. Berger A, Chandre F, Cornelie S, Paupy C. Controlling *Aedes* mosquitoes using densovirus-based biolarvicides: Current status and prospects. J Invertebr Pathol. 2025;211: 108314. doi:10.1016/j.jip.2025.108314

25. Perrin A, Gosselin-Grenet A-S, Rossignol M, Ginibre C, Scheid B, Lagneau C, et al. Variation in the susceptibility of urban *Aedes mosquitoes* infected with a densovirus. Sci Rep. 2020;10: 18654. doi:10.1038/s41598-020-75765-4

26. Drew GC, Stevens EJ, King KC. Microbial evolution and transitions along the parasite–mutualist continuum. Nat Rev Microbiol. 2021;19: 623–638. doi:10.1038/s41579-021-00550-7

27. Xu P, Chen F, Mannas JP, Feldman T, Sumner LW, Roossinck MJ. Virus infection improves drought tolerance. New Phytol. 2008;180: 911–921. doi:10.1111/j.1469-8137.2008.02627.x

28. Gorovits R, Shteinberg M, Anfoka G, Czosnek H. Exploiting Virus Infection to Protect Plants from Abiotic Stresses: Tomato Protection by a Begomovirus. Plants. 2022;11: 2944. doi:10.3390/plants11212944

29. Márquez LM, Redman RS, Rodriguez RJ, Roossinck MJ. A Virus in a Fungus in a Plant: Three-Way Symbiosis Required for Thermal Tolerance. Science. 2007;315: 513–515. doi:10.1126/science.1136237

30. Xu P, Liu Y, Graham RI, Wilson K, Wu K. Densovirus Is a Mutualistic Symbiont of a Global Crop Pest (*Helicoverpa armigera*) and Protects against a Baculovirus and Bt Biopesticide. PLOS Pathog. 2014;10: e1004490. doi:10.1371/journal.ppat.1004490

31. Jousset FX, Barreau C, Boublik Y, Cornet M. A parvo-like virus persistently infecting a C6/36 clone of *Aedes albopictus* mosquito cell line and pathogenic for *Aedes aegypti* larvae. Virus Res. 1993;29: 99–114. doi:10.1016/0168-1702(93)90052-o

32. Patsoula E, Samanidou-Voyadjoglou A, Spanakos G, Kremastinou J, Nasioulas G, Vakalis NC. Molecular and Morphological Characterization of *Aedes albopictus* in Northwestern Greece and Differentiation from *Aedes cretinus* and *Aedes aegypti*. J Med Entomol. 2006;43: 40–54. doi:10.1093/jmedent/43.1.40

33. Zhang Z, Reinikainen J, Adeleke KA, Pieterse ME, Groothuis-Oudshoorn CGM. Time-varying covariates and coefficients in Cox regression models. Ann Transl Med. 2018;6: 121. doi:10.21037/atm.2018.02.12

34. Rewitz KF, Yamanaka N, O’Connor MB. Chapter One - Developmental Checkpoints and Feedback Circuits Time Insect Maturation. In: Shi Y-B, editor. Current Topics in Developmental Biology. Academic Press; 2013. pp. 1–33. doi:10.1016/B978-0-12-385979-2.00001-0

35. De Moed GH, Kruitwagen CLJJ, De Jong G, Scharloo W. Critical weight for the induction of pupariation in *Drosophila melanogaster*: genetic and environmental variation. J Evol Biol. 1999;12: 852–858. doi:10.1046/j.1420-9101.1999.00103.x

36. Neckers L, Ivy SP. Heat shock protein 90. Curr Opin Oncol. 2003;15: 419. Available: https://journals.lww.com/co-oncology/abstract/2003/11000/heat_shock_protein_90.3.aspx

37. Thomas SR, Lengyel JA. Ecdysteroid-regulated heat-shock gene expression during *Drosophila melanogaster* development. Dev Biol. 1986;115: 434–438. doi:10.1016/0012-1606(86)90263-0

38. Briegel H, Timmermann SE. *Aedes albopictus* (Diptera: Culicidae): Physiological Aspects of Development and Reproduction. J Med Entomol. 2001;38: 566–571. doi:10.1603/0022-2585-38.4.566

39. Fischer K, Kölzow N, Höltje H, Karl I. Assay conditions in laboratory experiments: is the use of constant rather than fluctuating temperatures justified when investigating temperature-induced plasticity? Oecologia. 2011;166: 23–33. doi:10.1007/s00442-011-1917-0

